# Deconvolution of subcellular protrusion heterogeneity and the underlying actin regulator dynamics from live cell imaging

**DOI:** 10.1101/144238

**Authors:** Chuangqi Wang, Hee June Choi, Sung-Jin Kim, Aesha Desai, Namgyu Lee, Dohoon Kim, Yongho Bae, Kwonmoo Lee

## Abstract

Cell protrusion is morphodynamically heterogeneous at the subcellular level. However, the mechanistic understanding of protrusion activities is usually based on the ensemble average of actin regulator dynamics at the cellular or population levels. Here, we establish a machine learning-based computational framework called HACKS (deconvolution of Heterogeneous Activity Coordination in cytosKeleton at a Subcellular level) to deconvolve the subcellular heterogeneity of lamellipodial protrusion in migrating cells. HACKS quantitatively identifies distinct subcellular protrusion phenotypes from highly heterogeneous protrusion activities and reveals their underlying actin regulator dynamics at the leading edge. Furthermore, it can identify specific subcellular protrusion phenotypes susceptible to pharmacological perturbation and reveal how actin regulator dynamics are changed by the perturbation. Using our method, we discovered ‘accelerating’ protrusion phenotype in addition to ‘fluctuating’ and ‘periodic’ protrusions. Intriguingly, the accelerating protrusion was driven by the temporally coordinated actions between Arp2/3 and VASP: initiated by Arp2/3-mediated actin nucleation, and then accelerated by VASP-mediated actin elongation. We were able to confirm it by pharmacological perturbations using CK666 and Cytochalasin D, which specifically reduced ‘strong accelerating protrusion’ activities. Taken together, we have demonstrated that HACKS allows us to discover the fine differential coordination of molecular dynamics underlying subcellular protrusion heterogeneity via a machine learning analysis of live cell imaging data.

## Introduction

Cell protrusion is driven by spatiotemporally fluctuating actin assembly processes, and is highly morphodynamically heterogeneous at the subcellular level^1-3^. Elucidating the underlying molecular dynamics associated with subcellular protrusion heterogeneity is crucial to understanding the biology of cellular movement since protrusion determines the directionality and persistence of cell movements or facilitates the exploration of the surrounding environment^4^. Recent studies of the vital roles of cell protrusion in tissue regeneration^5,6^, cancer invasiveness and metastasis^7-9^, and the environmental exploration of leukocytes^10^ further emphasize the physiological and pathophysiological implication of understanding the fine molecular details of protrusion mechanisms. Although there has been considerable progress in analyzing individual functions of actin regulators, the precise understanding of how these actin regulators are spatiotemporally acting in cell protrusion is still limited. Moreover, it is a formidable task to dissect the actin regulator dynamics involved with cell protrusion because such dynamics are highly heterogeneous and fluctuate on both the micron length scale and the minute time scale^11-13^.

Although advances in computational image analysis on live cell movies have allowed us to study the dynamic aspects of molecular and cellular events at the subcellular level, the significant degree of heterogeneity in molecular and subcellular dynamics complicates the extraction of useful information from complex cellular behavior. The current method of characterizing molecular dynamics involves averaging molecular activities at the cellular level, which significantly conceals the fine differential subcellular coordination of dynamics among actin regulators. For example, our previous study employing the ensemble average method with the event registration showed that the leading edge dynamics of Arp2/3 and VASP were almost indistinguishable^13^. Over the past decade, hidden variable cellular phenotypes in heterogeneous cell populations have been uncovered by applying machine learning analyses^14,15^; however, these analyses primarily focused on static datasets acquired at the single-cell level, such as immunofluorescence^16^, mass cytometry^17^, and single-cell RNA-Seq^18^ datasets. Although some studies have examined the cellular heterogeneity of the migratory mode^19,20^, subcellular protrusion heterogeneity has not yet been addressed. Moreover, elucidating the molecular mechanisms that generate each subcellular phenotype has been experimentally limited because it is a challenging task to manipulate specific subclasses of molecules at the subcellular level with fine spatiotemporal resolution, even with the advent of optogenetics.

To address this challenge, we developed a machine learning-based computational analysis pipeline that we have called HACKS (deconvolution of Heterogeneous Activity Coordination in cytosKeleton at a Subcellular level) (Fig. 1) for live cell imaging data by an unsupervised machine learning approach combined with our local sampling and registration method^13^. HACKS allowed us to deconvolve the subcellular heterogeneity of protrusion phenotypes and statistically link them to the dynamics of actin regulators at the leading edge of migrating cells. Based on our method, we quantitatively identified subcellular protrusion phenotypes from highly heterogeneous and non-stationary edge dynamics of migrating epithelial cells. Furthermore, each protrusion phenotype was demonstrated to be associated with the specific temporal coordination of the actin regulators Arp2/3 and VASP at the leading edge. Our analysis of pharmacologically perturbed cells further demonstrated that the temporal coordination of Arp2/3 and VASP played an important role in driving accelerating protrusion. These results demonstrate that our machine learning framework HACKS provides a highly effective method to elucidate the fine differential molecular dynamics that underlie subcellular protrusion heterogeneity.

**Figure 1.**
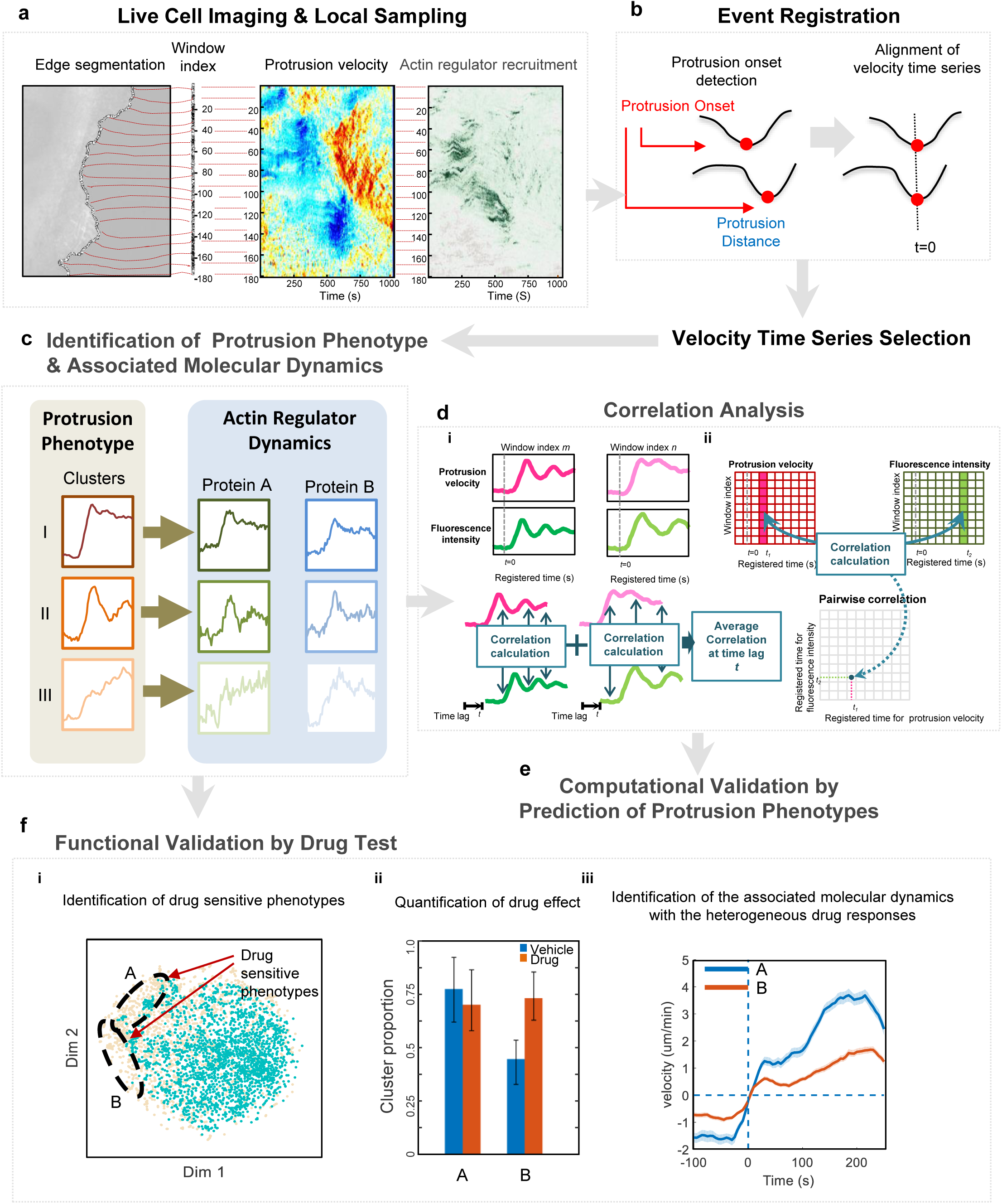
Schematic Representation of the Analytical Steps of HACKS. **(a)** Fluorescence time-lapse movies of the leading edges of a migrating PtK1 cell expressing Arp3-HaloTag were taken at 5 seconds/frame, and then probing windows (500 by 500 nm) were generated to track the cell edge movement and quantify protrusion velocities and fluorescence intensities. **(b)** The protrusion distance was registered with respect to protrusion onsets (t = 0). Time series of protrusion velocities were then aligned. **(c)** The protrusion phenotypes were identified with a time series clustering analysis, and associated with actin regulator dynamics. **(d, e)** Correlation/classification analysis between time series of the protrusion velocities and fluorescence intensities. Schematic diagrams of time lag (i) and time-specific correlation analysis (ii). **(f)** The hypotheses are validated functionally by molecular perturbation. The phenotypes susceptible to molecular perturbation are identified.

## Results

### A computational framework to deconvolve subcellular protrusion heterogeneity and the underlying molecular dynamics

To deconvolve the heterogeneity of the subcellular protrusion activity and their regulatory proteins at fine spatiotemporal resolution, we developed a computational analysis pipeline, HACKS (Fig. 1), which is based on an unsupervised machine learning method. HACKS allowed us to (1) identify distinct subcellular protrusion phenotypes based on a time series clustering analysis of heterogeneous subcellular protrusion velocities extracted from live cell movies (Fig. 1a-c), (2) associate each protrusion phenotype with pertinent actin regulator dynamics by comparing the average temporal patterns of protrusion velocities with those of actin regulators (Fig. 1c), (3) perform highly specified correlation and classification analyses of actin regulator dynamics of protrusion phenotypes to establish their association with fine mechanistic details (Fig. 1d-e), and (4) identify specific protrusion phenotypes susceptible to molecular perturbations, and functionally confirm the association between protrusion phenotype and the actin regulator dynamics (Fig. 1f). The framework can provide mechanistic insight into how the differential coordination of actin regulator dynamics organizes various subcellular protrusion phenotypes.

### Identification of subcellular protrusion phenotypes by a time series clustering analysis

Sample videos for the analysis were prepared by taking time-lapse movies of PtK1 epithelial cells expressing fluorescently tagged actin, Arp3, VASP and a cytoplasmic marker, HaloTag, with a spinning disk confocal microscope for approximately 200 frames at 5 sec/frame^11^ (Fig. 1a). Each time-lapse movie contains a single cell whose leading edge undergoes protrusion-retraction cycles. After segmenting the leading edge of each cell by multiple probing windows with a size of 500 by 500 nm^13^ (Fig. 1a), time series of velocities^11^ and fluorescence intensities of the tagged molecules^12,13^ acquired from each probing window were quantified (Fig. 1a). After registering protrusion onset at time zero (t=0), the time-series were aligned using the protrusion onset as a temporal fiduciary^13^ (Fig. 1b). To ensure a uniform time length of the data for the subsequent clustering analysis, we selected the first 50 frames (250 seconds) of protrusion segments, which is about the average protrusion duration^13^ from the pooled velocity time series. These protrusion segments were subjected to a further time series cluster analysis, whereas the fluorescence intensity data were later used to correlate the protrusion phenotypes and their underlying molecular dynamics.

The selected time series of the registered protrusion velocity contained a substantial amount of intrinsic fluctuations, hindering the identification of distinct clusters of similar protrusion activities. Therefore, we first denoised the time series velocity profile using Empirical Mode Decomposition^21^ and discretized the data using SAX (Symbolic Aggregate approximation, see Methods)^22^ to reduce the dimensionality and complexity of the data (Supplementary Text). We then extracted distinct patterns from fluctuating velocity time series by combining the autocorrelation distance measure with the Density Peak clustering^23^. The distance measures between different time series were calculated using the squared Euclidean distances between the corresponding autocorrelation functions (ACF) of each discretized time series. This autocorrelation distance partitioned the fluctuating time series of similar underlying patterns into the same clusters, enabling us to identify clusters with clear dynamic patterns (Supplementary Text). Following the ACF distance measure, we applied the Density Peak clustering algorithm, which has been shown to be superior to conventional K-means in partitioning data with complex cluster shapes^23^. As a result, the density-distance graph in Fig. 2p, where cluster centers are localized in the upper-right region (see Methods for detail), revealed five distinct clusters of subcellular protrusion activities. We also calculated three clustering criteria, Davies-Bouldin Index^24^, average silhouette value, and Calinski-Harabasz pseudo F-statistic^25^ as a function of the number of clusters, all of which suggested that the optimal number of clusters was five (Extended Data Fig. 3a-c). After the clustering analysis, average protrusion velocities and their 95% confidence intervals were calculated (Fig. 2h-m). Of note, after we tested different sets of algorithms, we found that ACF distance was the most important factor which allowed us to extract these distinct temporal patterns (Extended Data Fig. 1 and 2, Supplementary Text). Furthermore, we could not identify substantial differences among the velocity cluster profiles (dotted lines in Fig. 3g-ad) in each molecule (actin, Arp3, VASP, HaloTag), confirming that our clustering results are not skewed by a specific data set. The numbers of cells and probed windows used in the time series clustering analysis are presented in Extended Data Table 1.

**Figure 2.**
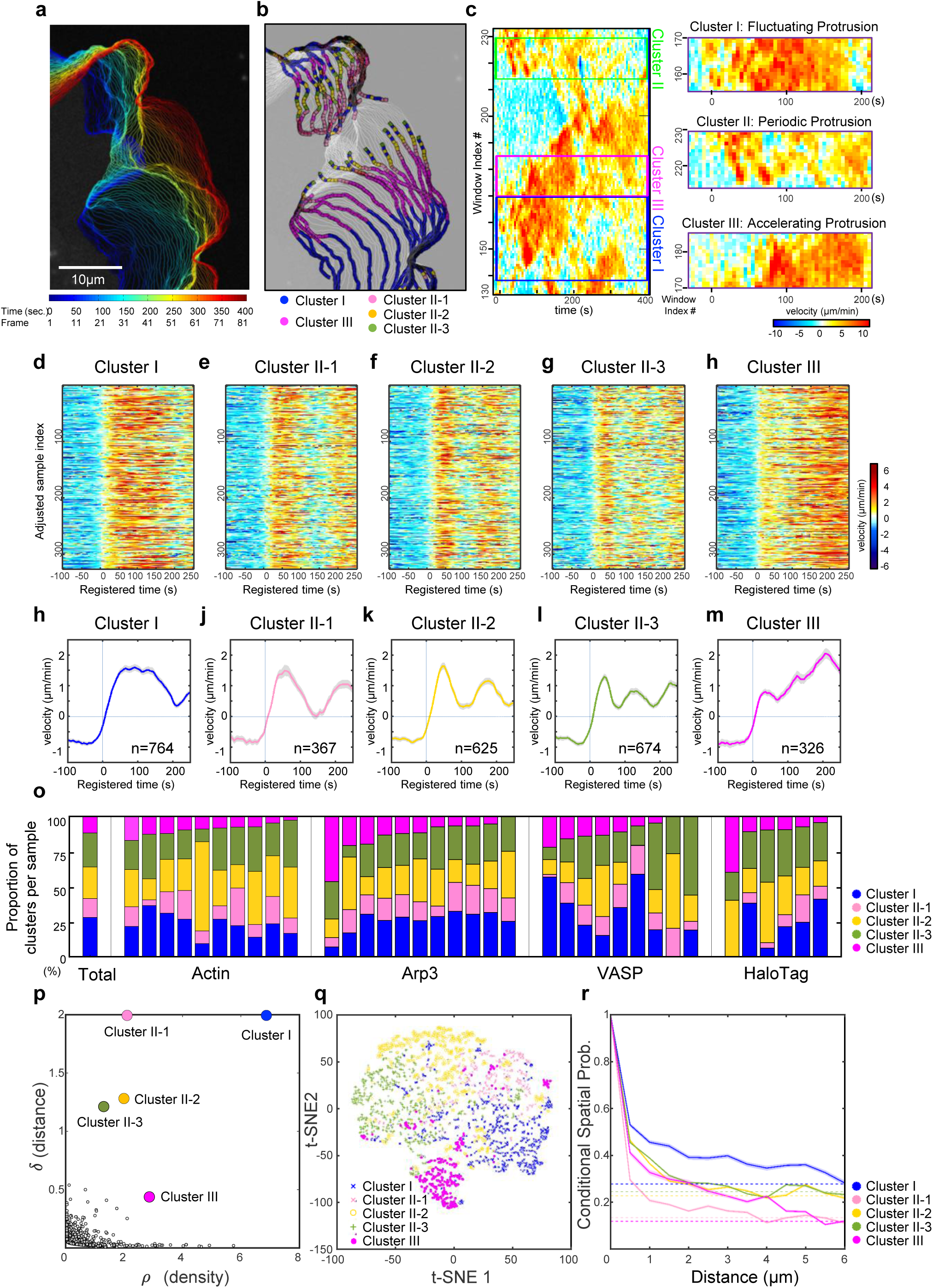
Subcellular Protrusion Phenotypes Revealed by a Time Series Clustering Analysis. **(a-c)** Representative examples of edge evolution (a), clustering assignments of each window every 10 frames (50 seconds) (b), and velocity activity maps (c) of a PtK1 cell stained with CellMask DeepRed. **(d-h)** Raw velocity maps for Cluster I, II-1, II-2, II-3, and III. **(h-m)** Average time series of protrusion velocity registered at protrusion onsets (t=0) in each cluster. Solid lines indicate population averages. Shaded error bands about the population averages indicate 95% confidence intervals of the mean computed by bootstrap sampling. n is the number of sampled time series. **(o)** Proportions of each cluster in total sample or individual cells expressing fluorescent actin, Arp3, VASP, and HaloTag, respectively. **(p)** Decision graph of the Density Peak clustering analysis of protrusion velocities. **(q)** t-SNE plot of the clusters. **(r)** Spatial conditional distribution of each cluster. Solid lines indicate population averages. Shaded error bands about the population averages indicate 95% confidence intervals of the mean computed by bootstrap sampling.

### Distinct subcellular protrusion phenotypes: the fluctuating, periodic and accelerating protrusion

The visual inspection of the average velocity profiles of the identified clusters (Fig. 2h-m) demonstrated that the overall differences among the protrusion phenotypes originated from differences in the timing and number of peaks the velocity reached. Whereas Cluster I did not exhibit dramatic changes in protrusion velocities after reaching its peak at the earlier part of the protrusion segment (Fig. 2h), the remaining clusters exhibited substantial acceleration or deceleration in the protrusion velocities with varying timing and number. Clusters II-1 (Fig. 2j), II-2 (Fig. 2k), and II-3 (Fig. 2l) exhibited periodic fluctuation in the acceleration and deceleration of protrusion with differential timing, reaching their velocity peak more than twice. Conversely, Cluster III (Fig. 2m) demonstrated persistently accelerating behavior where protrusion velocities continued to increase until the late phase of the protrusion. Clusters I, II-1, II-2, II-3 and III comprised 27.7%, 13.3%, 22.7%, 24.5%, and 11.8% of the entire sample, respectively, and individual cells expressing different fluorescent proteins exhibited similar tendencies (Fig. 2o, Extended Data Fig. 3h), suggesting the intracellular origin of protrusion heterogeneity. Nevertheless, cell-to-cell variability in cluster distribution persisted, suggesting that the clusters may also reflect individual cellular responses to differential cellular contexts or microenvironments.

The validity of our clustering result was confirmed by visually inspecting the velocity activity map (Fig. 2d-h). Clusters II-1/2/3 (Fig. 2e-g) and III (Fig. 2h) exhibited clearly distinguishable patterns, whereas Cluster I (Fig. 2f) contained fluctuating velocity profiles (See Extended Data Fig. 3g for the full maps). The t-SNE (Fig. 2r), multidimensional scaling, silhouette, and order distance plots (Extended Data Fig. 3d-f) of the clustering results further confirmed the stability and tightness of Clusters II-1/2/3 and III but suggested residual heterogeneity in Cluster I, which is in agreement with the velocity activity maps (Fig. 1d). To quantify the spatial structure of the protrusion phenotypic clusters, we estimated the conditional probability that the same cluster exists over the distance from a given cluster (Fig. 2r). As the distance increases between two neighboring clusters, this conditional probability in all clusters decreases to their basal levels of the cluster proportions (Fig. 2o). The conditional probability in Cluster II-1/2/3 quickly decreased within 2 μm distance whereas those in Cluster I and III persisted up to 5 μm (Fig. 2r). This data suggests that Clusters I and III aggregate and act collectively more so compared to Cluster II-1/2/3. In addition to PtK1 cells, we further performed the same analysis on MCF10A, human mammary epithelial cells. MCF10A also had very similar subcellular protrusion patterns (Extended Data Fig. 4), suggesting that these results by HACKS are not limited to a specific cell line.

Notably, this procedure revealed the differential subcellular protrusion phenotypes with distinct velocity profiles using our time series clustering framework. These variabilities were not previously elucidated when the entire time series was ensemble averaged without taking into account the subcellular protrusion heterogeneity^13^. The visualization of the edge evolution (Fig. 2a), the cluster assignments evolution (Fig. 2b), and the protrusion velocity map (Fig. 2c) of the exemplified live cell movie representatively manifested the morphodynamic features of each subcellular protrusion phenotype (Supplementary Movie 1). Since Cluster II clearly exhibited periodic edge evolution, we refer to Cluster II-1/2/3 collectively as ‘periodic protrusion.’ Conversely, Cluster III showed accelerated edge evolution, and, therefore, we refer to Cluster III as ‘accelerating protrusion.’ As summarized in Extended Data Table 2, this phenotype was found by the specific combination of SAX and ACF dissimilarity measure (Supplementary Text). To the best of our knowledge, this study is the first to quantitatively characterize persistently ‘accelerating’ cell protrusion. Previous studies described persistent protrusion based on the protrusion distance over longer time scales^11,13,26,27^, but have not focused on ‘accelerating’ protrusion. Finally, Cluster I was named ‘the fluctuating protrusion’ cluster because of the irregularity of its velocity profiles.

### Differential molecular dynamics of actin regulators underlying the accelerating protrusion phenotype

We hypothesized that the distinctive subcellular protrusion phenotypes arise from the differential spatiotemporal regulation of actin regulators. Therefore, we next investigated the relationship between the velocity profiles of each protrusion phenotype and the fluctuation of the signal intensities of actin and several actin regulators for each protrusion phenotype. We selected a set of fluorescently tagged molecules to be expressed and monitored; SNAP-tag-actin, HaloTag-Arp3 (tagged on the C-terminus), which represented the Arp2/3 complex involved in actin nucleation, and HaloTag-VASP or GFP-VASP, which represented actin elongation. A diffuse fluorescent marker, HaloTag labeled with tetramethylrhodamine (TMR) ligands^28^, was used as a control signal. The fluorescence intensities of each tagged molecule were acquired from each probing window along with the protrusion velocities (Fig. 1a). The time-series of the fluorescence intensities of each molecule were then grouped and averaged according to the assigned protrusion phenotype (Fig. 1c and Fig. 3g-ad).

**Figure 3.**
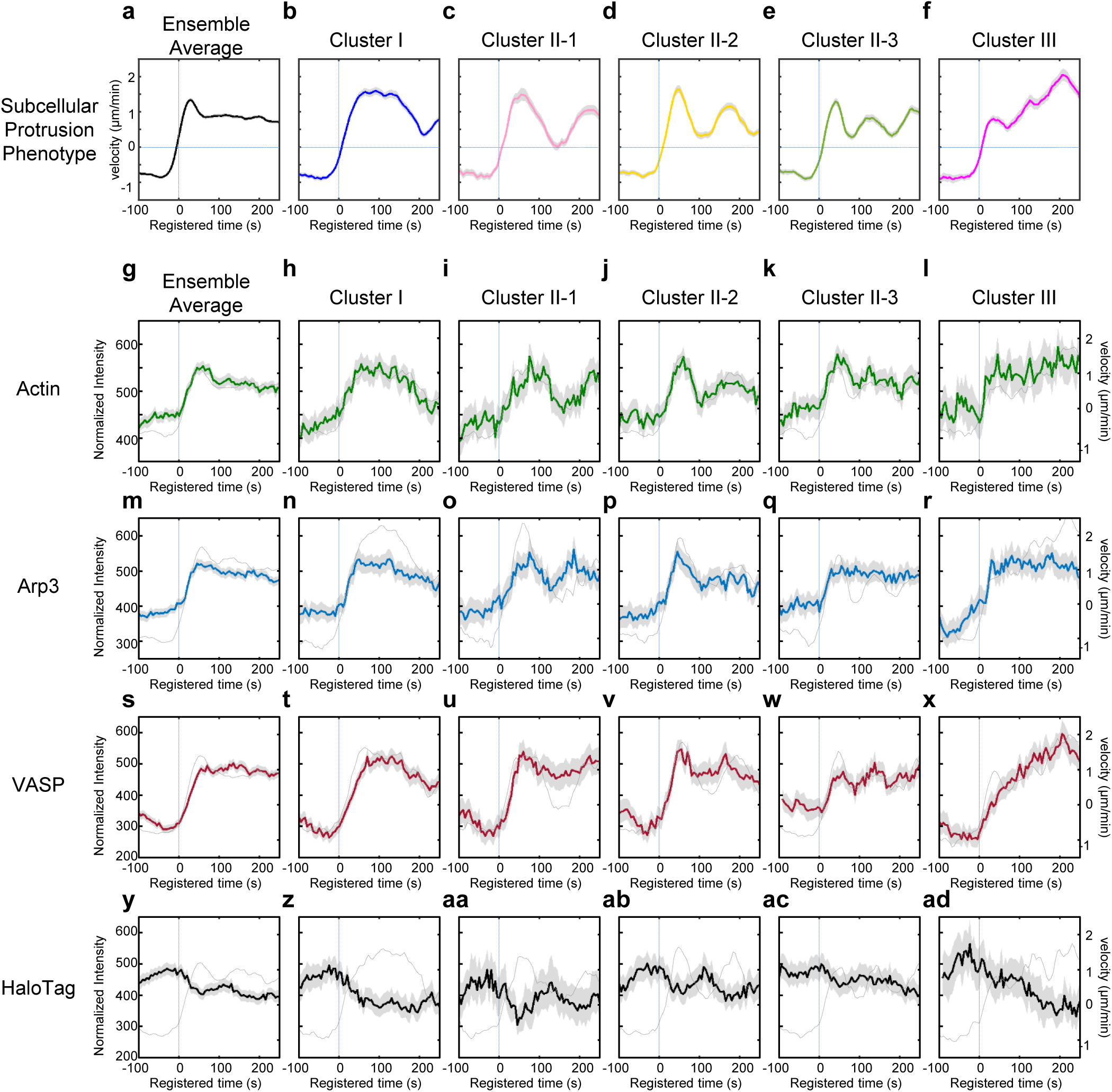
Identified Distinctive Subcellular Protrusion Phenotypes and Their Differential Underlying Molecular Dynamics. **(a-ad)** Protrusion velocity and normalized fluorescence intensity time series registered with respect to protrusion onset for ensemble averages of entire samples, Cluster I, Cluster II-1, Cluster II-2, Cluster II-3, and Cluster III. Solid lines indicate population averages. Shaded error bands about the population averages indicate 95% confidence intervals of the mean computed by bootstrap sampling. The dotted lines in (g-ad) indicate protrusion velocity time series associated with the specified fluorescent proteins.

Whereas the molecular dynamics of actin, Arp3 and VASP all exhibited patterns similar to those of the velocity profiles in Clusters I and II-1 (Fig. 3b-c, h-i, n-o, t-u,T), the Arp3 temporal patterns became less correlated with those of protrusion velocity in Cluster II-2 and II-3 as the frequency of the oscillation increased (Fig. 3p-q). This demonstrates that underlying molecular temporal patterns can be highly variable depending on dynamic properties of protrusion activities. Intriguingly, Cluster III also exhibited distinctive differential molecular dynamics among the molecules and in relation to velocity profiles (Fig. 3f, l, r, x). Specifically, whereas the protrusion velocity continued to increase until the late stages of the protrusion segment in the accelerating protrusion (Cluster III) (Fig. 3f), the actin fluorescence intensity soon reached its maximum in the early phase and remained constant (Fig. 3l). This pattern indicates that edge movement during accelerating protrusion is mediated by the elongation of existing actin filaments rather than *de novo* actin nucleation. Conversely, Clusters I and II-1/2/3 exhibited increased actin intensity at the leading edge along with increased protrusion velocity (Fig. 3h-k), indicating that actin nucleation mediates subcellular protrusion.

In accordance with the plateaued actin intensities in Cluster III (Fig. 3l), the Arp3 intensity remained constant after reaching its peak in the early protrusion phase (Fig. 3r), whereas the VASP intensities began to increase at protrusion onset and continued to increase (Fig. 3x). These findings suggest that actin elongation by VASP plays a crucial role in driving accelerating protrusion. Whereas the Arp2/3 complex has been considered as a major actin nucleator that drives lamellipodial protrusion^29^, Arp2/3 seemed to play a role in the earlier part of the protrusion in accelerating protrusion. Approximately 50 seconds after protrusion onset, the Arp3 intensity reached its peak (Fig. 3r), and the acceleration temporarily stopped (Fig. 3f). Notably, the Arp3 intensities began to increase 50 seconds prior to the protrusion onset in Cluster III (Fig. 3r), whereas they began to increase at the onset of the protrusion in Clusters I and II-1/2/3 (Fig. 3n-q). These findings imply that there exists specific temporal coordination where the Arp2/3 complex nucleates actin networks in the early phase, and VASP then elongates actin filaments to drive the later stages of accelerating protrusion. The specificity of the relationship between the protrusion phenotypes and the underlying molecular dynamics was further validated with a control experiment using HaloTag-TMR (Fig. 3y-ad). Diffused cytoplasmic fluorescence did not exhibit any cluster-specific pattern. Instead, it inversely correlated with the protrusion velocity, suggesting that the cell edges become thinner as the protrusion velocity increases^13^. Notably, the differential dynamics of Arp3 and VASP were not observed when the entire time series dataset was ensemble averaged^13^ (Fig. 3m and s). These results demonstrate the power of our computational framework in revealing the hidden differential subcellular dynamics of actin regulators involved in the generation of heterogeneous morphodynamic phenotypes.

### VASP recruitment strongly correlates with protrusion velocity

To quantitatively assess the coordination between protrusion velocities and the dynamics of actin regulators, we performed a time-lag correlation analysis by calculating Pearson’s correlation coefficients between protrusion velocities and actin regulator intensities with varying time lags in the same windows and averaged over different sampling windows (Fig. 1d.i). For actin and Arp3, the significant but relatively weak correlations were identified between the protrusion velocity and the intensities in all clusters (Fig. 4a-b). Conversely, the correlation of VASP in all clusters was stronger, particularly the correlation in Cluster III being the strongest in all clusters (Fig. 4c). Consistent with the results of cytoplasmic dynamics (Fig. 3y-ad), HaloTag-TMR intensities were negatively correlated with protrusion velocities (Fig. 4d). Furthermore, a comparison of the maximum correlations in each cluster showed that VASP exhibited significantly stronger correlations than the Arp2/3 complex in all clusters (Fig. 4e, Extended Data Table 3). These findings suggest that VASP may play a more direct role in mediating protrusion velocities in all clusters than Arp2/3.

**Figure 4.**
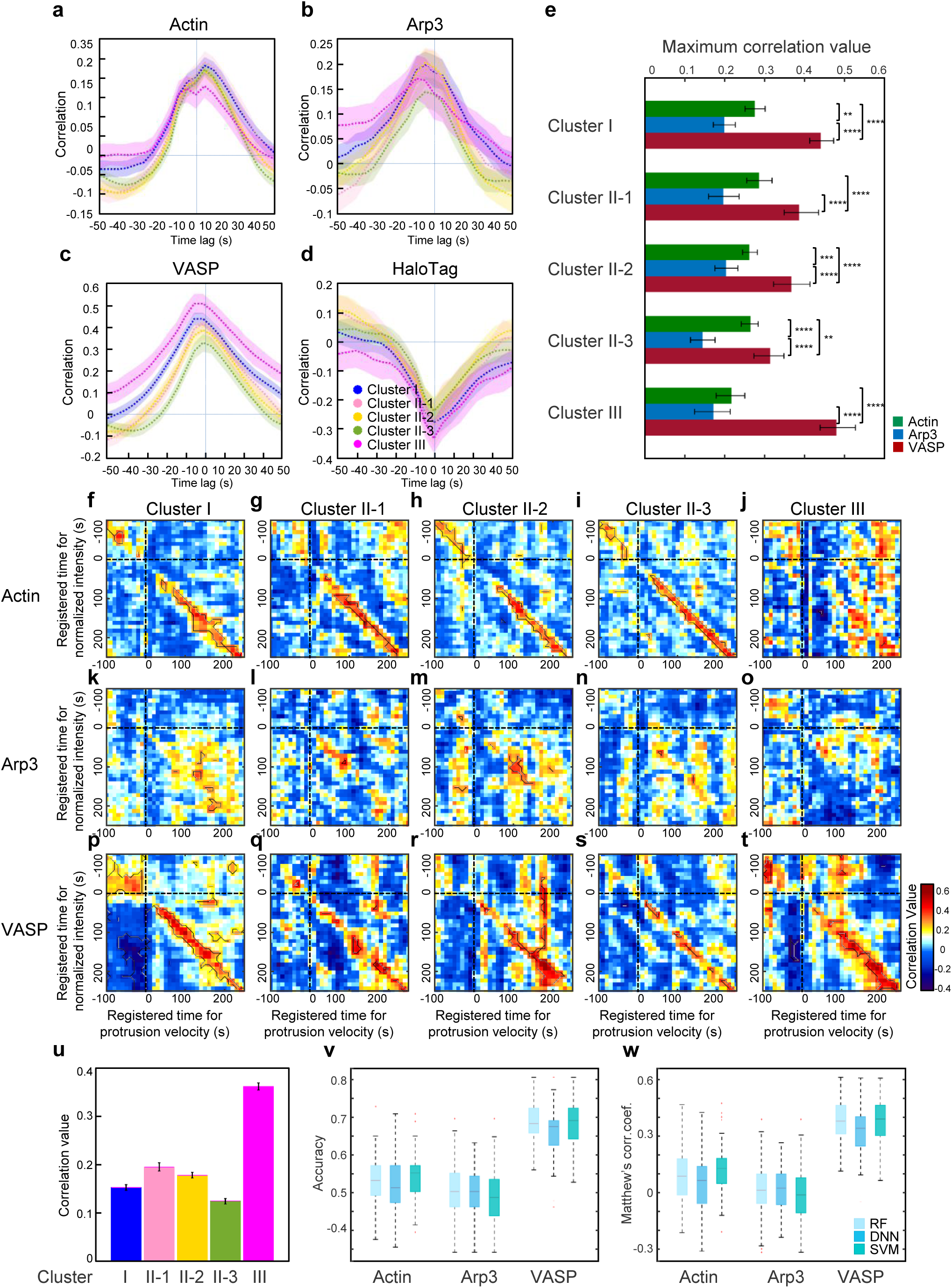
Correlation and Classification Analyses between Protrusion Velocity and Actin Regulator Dynamics. **(a-d)** Pearson’s cross-correlation of edge velocity and actin (a), Arp3 (b), VASP (c), and HaloTag (d) as a function of the time lag between two time series. Solid lines indicate population averages. Shaded error bands about the population averages indicate 95% confidence intervals of the mean computed by bootstrap sampling. **(e)** Comparison and statistical testing of maximum correlation coefficients from (a-d) in each cluster. ** (p < 0.01), *** (p < 0.001) and **** (p < 0.0001) indicate the statistical significance by two-sample Kolmogorov-Smirnov (KS) test. The p-values are listed in Extended Data Table 3. **(f-t)** Pairwise Pearson’s correlation coefficients of protrusion velocity and fluorescence intensity time series registered relative to protrusion onset. The regions surrounded by the black lines are statistically significant correlation by Benjamini-Hochberg multiple hypothesis testing. **(u)** Pearson’s correlation coefficients between early Arp3 intensities and late protrusion velocities in each cluster. The error bar indicates 95% confidence interval of the mean by bootstrapping. **(v-w)** Accuracy (v) and Matthews correlation coefficients (w) of the classification analysis of Cluster III against Clusters I/II.

Although the above-described conventional time correlation analysis effectively demonstrated the overall correlation between molecular dynamics and the protrusion velocity, its ability to reveal changes in this correlation over time as the protrusion progresses is limited. In other words, the correlation between the protrusion velocities and the fluorescence intensities for each specific time point was not examined in the previous analyses (Fig. 4a-d). Therefore, we performed sample-based correlation analyses whereby calculating pairwise Pearson correlation coefficients, c({*V*}_*t_j_*_, {*I*}_*t_j_*_), between the sample of the protrusion velocity, {*V*}_*t_j_*_, at the registered time, *t*_*i*_, and the sample of the actin regulator intensity {*I*}_*t_i_*_, at the registered time, *t*_*j*_, over the entire probing window population (Fig. 1d.ii)^13^. Then, the statistical significance of the correlations were tested by Benjamini-Hochberg multiple testing^30^.

As expected, the pairwise time correlation analysis between the actin intensities and protrusion velocities (Fig. 4f-j) further supported the proposition that accelerating protrusions are mediated by the elongation of pre-existing actin filaments, whereas actin nucleation is responsible for non-accelerating protrusions. The significant regions (the black boundaries in Fig. 4f-t) of instantaneous positive correlations between the actin intensities and protrusion velocities at the leading edge found in Clusters I and II-1/2/3 (Fig. 4f-i) were absent in Cluster III (Fig. 4j). Notably, in the previous time lag correlation analysis, the weak correlation for actin persisted in Cluster III (Fig. 4a). This finding suggests that pairwise correlations at specific time points can effectively and more precisely reveal the various aspects of the coordination between protrusion velocities and the underlying molecular dynamics.

Intriguingly, we did not identify a similarly significant instantaneous correlation between the protrusion velocity and Arp3 in any cluster (Fig. 4k-o). Conversely, we identified a significantly stronger instantaneous correlation between the VASP intensities and protrusion velocities in all clusters in the time-specific correlation analysis (Fig. 4p-t). This is consistent with the previous study such that the edge velocity and lamellipodial VASP intensity were highly correlated when the leading edges of B16 melanoma cells had a uniform rate of protrusion^31^; but our study provided substantial quantitative evidence from the samples exhibiting highly heterogeneous and non-stationary edge movements. This further suggests that VASP compared to Arp2/3 plays a more direct role in controlling the protrusion velocity at the leading edge in all protrusion clusters. In Cluster I and II-1/2/3, VASP-dependent actin elongation likely tightly coordinates with Arp2/3 complex-mediated actin nucleation because actin exhibited a strong instantaneous correlation with protrusion velocity. Conversely, the significant and strong instantaneous correlation between VASP and the protrusion velocity, which begins to appear 100 seconds after protrusion onset (Fig. 4t), along with no correlation between actin and the protrusion velocity in Cluster III (Fig. 4j) suggests that actin elongation by VASP plays a key role in accelerating protrusion. Since the Arp3 intensity in Cluster III reached the maximum at approximately this time point (Fig. 3r), we hypothesized that Arp2/3 in the early phase of Cluster III might play an important role. To test this idea, we integrated early Arp3 intensities between 0 to 50 seconds and correlated them with average protrusion velocities between 150 and 200 seconds in each cluster (Fig. 4u, Extended Data Fig. 5c). The Pearson’s correlation coefficient Cluster III was significant (0.36, p = 0.0002) and larger than those in other clusters. This is consistent with our hypothesis that Arp2/3 may be important in initiating cell protrusion in Cluster III.

Notably, both the strong correlation between VASP and the protrusion velocity observed in all clusters and the postulated mode of VASP in regulating accelerating protrusions suggest that VASP plays a more critical role in generating differential protrusion phenotypes. The differences in how VASP and Arp2/3 polymerize actin further validate our interpretation. VASP facilitates actin filament elongation by binding to the barbed ends of actin filaments at the leading edge^32-34^, whereas Arp2/3 binds to the sides of the mother filaments and initiates actin nucleation. Thus, the ability of Arp2/3 to directly control bared end elongation is limited^35^. Because actin elongation at the barbed end pushes the plasma membrane and generates protrusion velocity, the strong correlation between VASP activity and protrusion velocity at the leading edge is plausible.

Since VASP intensities were well correlated with protrusion velocities, we next investigated whether VASP intensities contain sufficient information to predict protrusion phenotypes. To this end, we applied supervised learning approaches to further validate that VASP intensity time series can predict the acceleration phenotype. Using support vector machine (SVM), deep neural network (DNN), and random forest (RF), we built classifiers from the normalized intensities of actin, Arp3, and VASP to distinguish the non-accelerating (Clusters I and II-1/2/3) and accelerating (Cluster III) protrusion phenotypes. As a result, classifiers that were trained using VASP intensities could distinguish accelerating protrusions from non-accelerating protrusions with a significantly higher accuracy (Fig. 4v) and MCC (Matthews correlation coefficient) (Fig. 4w) than the classifiers trained using the actin and Arp3 intensities (p-values in Extended Data Table 4). Despite the modest classification accuracy, this difference suggests not only that VASP correlated with protrusion velocity but also that the correlation is strong enough to predict the accelerating protrusion phenotype.

### Deconvolution of heterogeneity of protrusion responses to pharmacological perturbations at the subcellular level

Our statistical analyses thus far suggest that the early recruitment of Arp2/3 at the leading edge leads to VASP recruitment to barbed ends of actin filaments, giving rise to accelerating cell protrusion. Since Arp2/3 was implicated in the early phase of accelerating protrusion, we treated PtK1 cells with an Arp2/3-specific inhibitor, CK666^36^ (50 μM) to validate the functional role of Arp2/3. Remarkably, CK666 treated cells still exhibited highly active protrusion activities with 50 μM concentration, and they were visually indistinguishable from the control cells treated with the inactive compound, CK689. After pooling CK666 and CK689 data together, we performed the time series clustering analysis. CK666 and CK689 treated cells still exhibited similar temporal patterns in all clusters (Fig.5f-k, Extended Data Fig. 6, Supplementary Movie 2 and 3), even if the protrusion velocities in Cluster I, II-1, and III were modestly reduced by CK666 (Fig. 5g, h, k). The t-SNE visualization of the autocorrelation functions (ACFs) of all protrusion time series revealed that CK666 (Fig. 5b) affected two densely populated areas in the control (CK689) cells (the dotted circles in Fig. 5a-b), and overlaying the cluster assignment in these t-SNE plots revealed that Cluster III was reduced by CK666 (Fig. 5c-d). The quantification of the proportion of each cluster confirmed that Cluster III was significantly reduced by the CK666 treatment (Fig. 5e, p = 0.0059). In turn, this led to the significant increase of Cluster II-1 (Fig. 5e, p = 0.0001). Intriguingly, the other clusters were not significantly affected by CK666, suggesting that the reduced Arp2/3 activities could be compensated by other actin regulators^37^. These results verify that Arp2/3 plays a specific functional role in accelerating protrusion. Furthermore, these demonstrate that our HACKS framework enables us to identify the susceptible clusters, which respond specifically to pharmacological perturbations.

**Figure 5.**
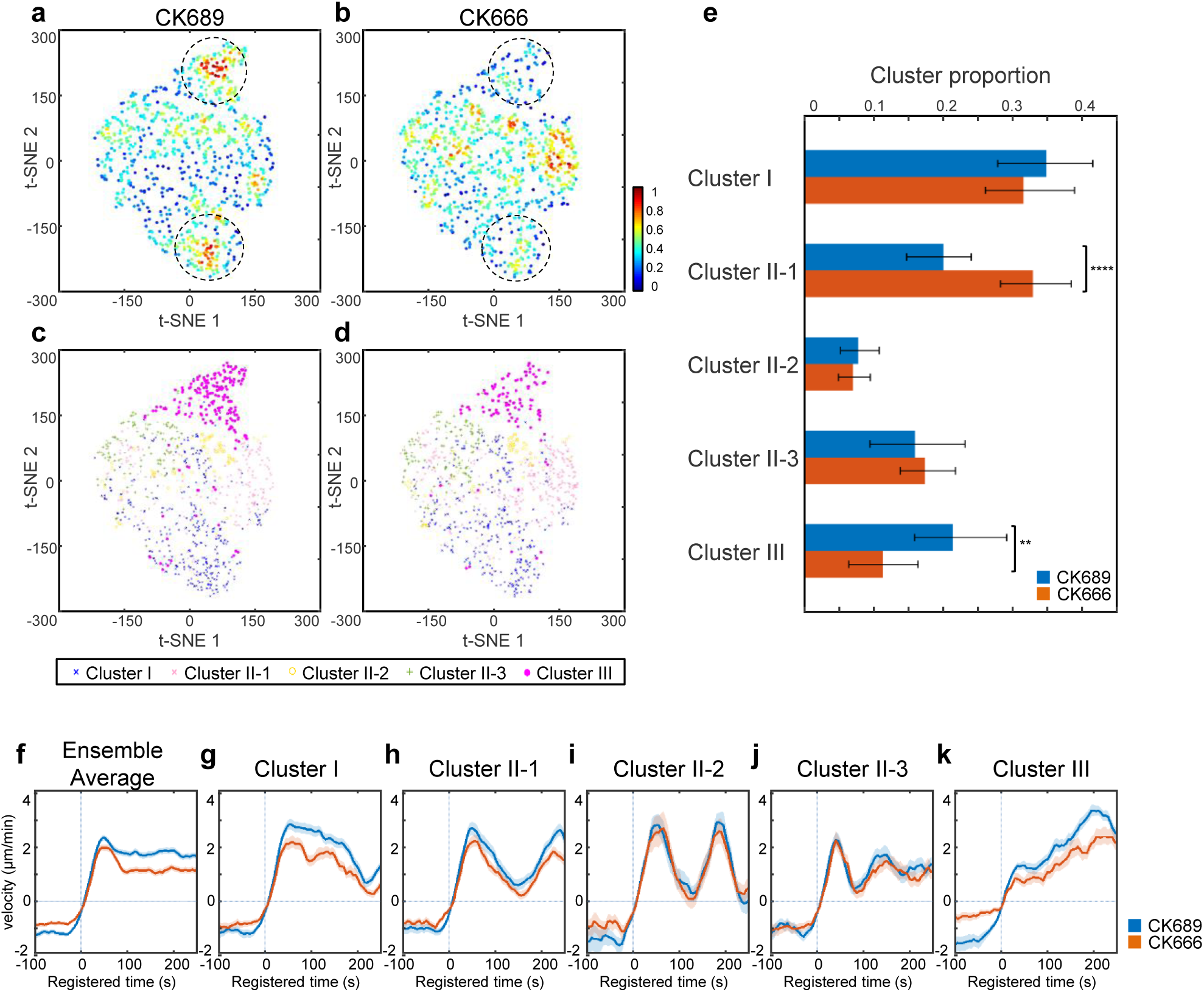
Responses of subcellular protrusion phenotypes to Arp2/3 inhibitor, CK666. **(a-d)** t-SNE plots of autocorrelation functions of protrusion velocity time series overlaid with the density of data (a-b) and cluster assignments (c-d). **(e)** Comparison of the proportions of each cluster between CK689 and CK666. Error bars indicate 95% confidence interval of the mean of the cluster proportions. ** (p < 0.01) and **** (p < 0.0001) indicates the statistical significance by bootstrap sampling. **(f-k)** Protrusion velocity time series registered with respect to protrusion onset for ensemble averages of the entire sample set, and each cluster in CK689 and CK666 treated cells. Solid lines indicate population averages. Shaded error bands about the population averages indicate 95% confidence intervals of the mean computed by bootstrap sampling.

Next, to validate the functional role of VASP in accelerating protrusion (Cluster III), we treated PtK1 cells with low concentrations (50 and 100 nM) of Cytochalasin D (CyD) to displace VASP from the barbed ends of actin filaments^26,38-40^. Using immunofluorescence, we confirmed that the CyD treatment effectively removed the phosphorylated VASP, which is a functional form of VASP, from the lamellipodial leading edge of PtK1 cells (Extended Data Fig. 7). Consistent with our previous correlation analyses where VASP intensities correlated with protrusion velocities in all clusters, the time series clustering analysis using the pooled DMSO and CyD (50, 100nM) data revealed that protrusion velocities in all protrusion clusters in the CyD treated cells were significantly reduced in a dose-dependent manner in comparison to DMSO treated cells (Fig. 6h-m, Extended Data Fig. 8, Supplementary Movie 4-6). Nonetheless, the CyD treated cells retained similar clustering structures, demonstrating the specificity of the CyD treatment in these low concentrations. The t-SNE plots of ACFs of each velocity time series also revealed that two dense areas were affected by the CyD treatment (the dotted circles in Fig. 6a-c), which includes the region of Cluster III (Fig. 6d-f). The proportion of Cluster III was significantly but modestly reduced by the CyD treatment (Fig. 6g, p = 0.043 for 50 nM, 0.018 for 100 nM).

**Figure 6.**
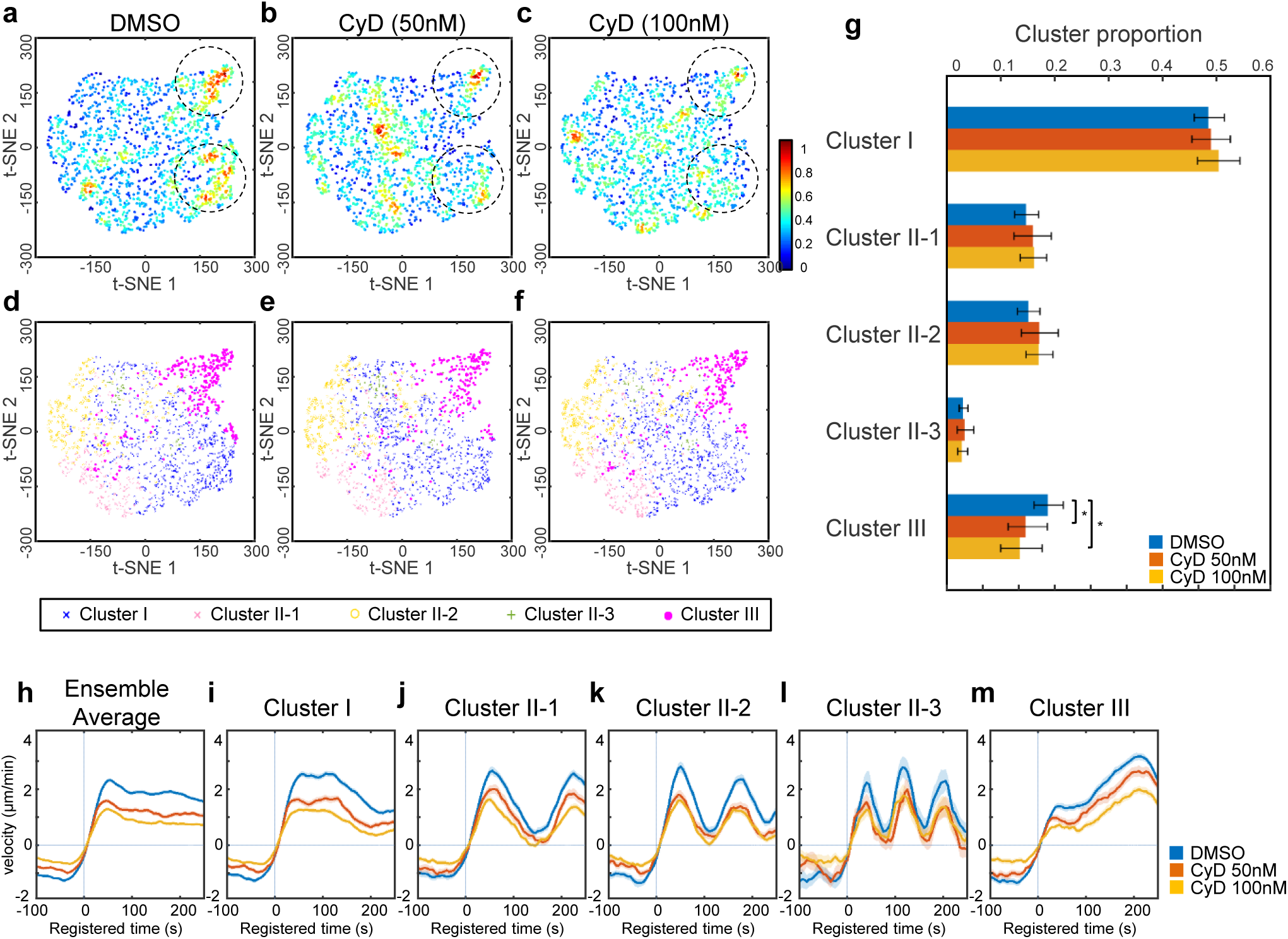
Responses of subcellular protrusion phenotypes to low doses of Cytochalasin D. **(a-f)** t-SNE plots of autocorrelation functions of protrusion velocity time series overlaid with the density of data (a-c) and cluster assignments (d-f). **(g)** Dose-response of the proportions of clusters to CyD. Error bars indicate 95% confidence interval of the mean of the cluster proportions. * (p < 0.05) indicates the statistical significance by bootstrap sampling. **(h-m)** Protrusion velocity time series registered with respect to protrusion onset for ensemble averages of entire samples, each cluster in DMSO and CyD treated cells. Solid lines indicate population averages. Shaded error bands about the population averages indicate 95% confidence intervals of the mean computed by bootstrap sampling.

We observed CyD treatment tended to reduce the overall protrusion velocities. Therefore, we visualized the data distributions using t-SNE with denoised protrusion velocities instead of ACFs to further investigate the effects of CyD on Cluster III in terms of regulation of protrusion velocity. This t-SNE analysis revealed high-density regions of the subcellular protrusion velocities which are highly susceptible to the CyD and CK666 treatment (the dotted circles in Fig. 7a-e). Overlaying the cluster assignments in these t-SNE plots showed that Cluster III contained a substantial portion of the CyD and CK666-susceptible regions (Extended Data Fig. 9a). Intriguingly, the tSNE plots of Cluster III of the control cells (Extended Data Fig. 9b) suggest that Cluster III can be largely grouped into two, which may have differential susceptibilities to CyD and CK666. Therefore, we further divided Cluster III into two sub-clusters (Fig. 7f-j) based on denoised protrusion velocities pooled from CyD and CK666 dataset by a community detection algorithm^41^ (Extended Data Fig. 9c-e). While both Cluster III-1 and III-2 (Fig. 7f-j) maintained similar temporal patterns, Cluster III-2 had substantially stronger accelerating activities compared to Cluster III-1 (DMSO in Fig. 7m-n; CK689 in Fig. 7o-p). Intriguingly, the t-SNE plots revealed that ‘strongly accelerating protrusion’ (Cluster III-2) was preferentially affected by the CyD (Fig. 7f-h) and CK666 (Fig. 7i-j) treatment. The quantification of the proportion of these sub-clusters (Fig. 7k) confirmed that strongly accelerating protrusion (Cluster III-2) was significantly reduced by the CyD treatment in comparison to DMSO treatment in a dose-dependent manner (p = 0.024 for 50 nM, < 0.0001 for 100 nM), whereas the weakly accelerating protrusion (Cluster III-1) was increased (p = 0.006 for 100 nM). Therefore, the average protrusion velocities in Cluster III in CyD treatment were significantly reduced to be comparable to Cluster III-1 in DMSO treatment and was significantly lower than Cluster III-2(Fig. 7m-n). Consistently, the proportion of strongly accelerating protrusion (Cluster III-2) was significantly reduced by the CK666 treatment (Fig. 7l, p = 0.0026) and the average velocities of Cluster III in CK666 treatment were also reduced to those of weakly accelerating protrusion (Cluster III-1) in CK689 treatment. These data demonstrate HACKS allowed us to successfully identify the drug-susceptible sub-phenotypes, where strongly accelerating protrusion is specifically affected by Arp2/3 and VASP inhibition.

**Figure 7.**
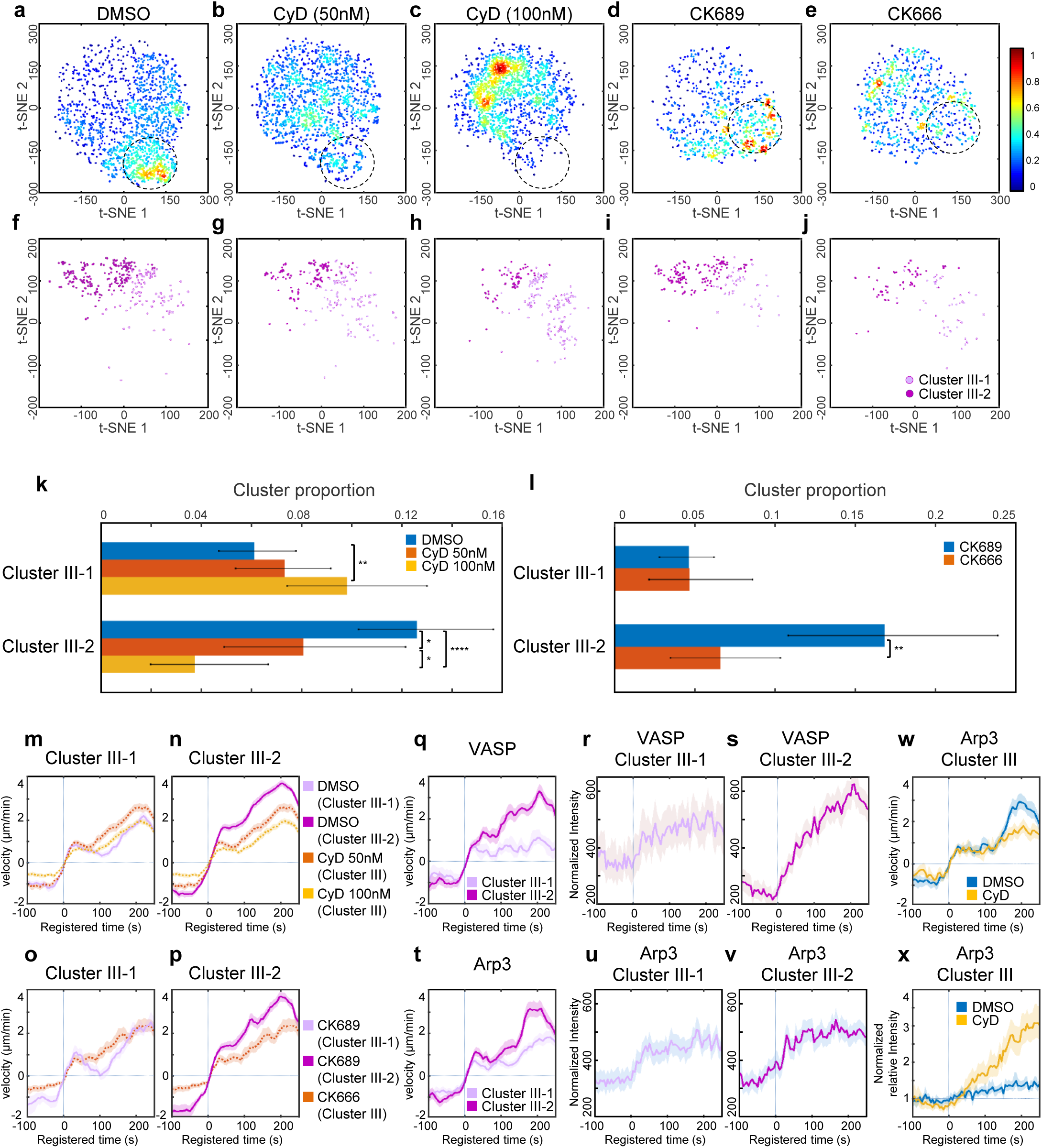
Effects of Cytochalasin D and CK666 on strongly accelerating protrusion. **(a-e)** t-SNE plots of the denoised protrusion velocity time series of the whole sample overlaid with the density of data. **(f-j)** t-SNE plots of the denoised velocities of the sub-clusters (Cluster III-1 and III-2) in Cluster III. **(k)** Response of the proportions of clusters to CyD (k) and CK666 (l) treatment. Error bars indicate 95% confidence interval of the mean of the cluster proportions. * (p < 0.05), ** (p < 0.01), and **** (p < 0.0001) indicate the statistical significance by bootstrap sampling. **(m-p)** Protrusion velocity time series registered with respect to protrusion onset for ensemble averages of Cluster III-1 and III-2 in CyD (m-n) and CK666 (o-p) treated cells. **(q-v)** Protrusion velocity and normalized fluorescence intensity time series of VASP (q-s) and Arp3 (tv) registered with respect to protrusion onset for ensemble averages of Cluster III-1 and III-2 in unperturbed PtK1 cells. **(w-x)** Protrusion velocity (w) and normalized fluorescence intensity time series of Arp3 (x) registered with respect to protrusion onset for ensemble averages of Cluster III in DMSO and 100 nM CyD treated cells. Solid lines indicate population averages. Shaded error bands about the population averages indicate 95% confidence intervals of the mean computed by bootstrap sampling.

Next, we further investigated whether dynamics of VASP and Arp3 in accelerating protrusion is differentially regulated between Cluster III-1 and Cluster III-2. We divided the intensity time series of VASP and Arp3 in Cluster III (Fig. 3r and x) into two sub-clusters and compared their differential dynamics. The recruitment dynamics of VASP in Cluster III-2 exhibited strong increase, while that of Cluster III-1 exhibited only moderate elevation, which is within the 95% confidence interval of the mean (Fig. 7r). On the other hand, Arp3 intensity patterns in Cluster III-1 and 2 were almost identical (Fig. 7u-v). This is consistent with our notion that Arp2/3 is involved in initiating accelerating protrusion and VASP is important in the output of accelerating protrusion. To functionally confirm this, we compared Arp3-GFP fluorescence dynamics at the leading edges in each cluster without and with 100 nM CyD treatment (Extended Data Fig. 10). To this end, we normalized Arp3 intensities at the leading edge by those of the lamella region in the same cell to quantitatively compare the Arp3 accumulation in different experimental condition. Under CyD treatment, the Arp3 fluorescence normalized by lamella intensity still started to increase at the protrusion onset in Cluster III (Fig. 7x). Normalized Arp3 fluorescence continued to increase up to 4 fold more than the DMSO control while the protrusion velocity did not increase (Fig. 7w). First, this suggests that CyD treatment did not affect the initial Arp2/3 recruitment to the leading edge in accelerating protrusion, which proposes that Arp2/3 precedes VASP in accelerating protrusion. In addition, this data shows that even increasing Arp2/3 recruitment under CyD treatment could not produce strongly accelerating protrusion without VASP activity. Therefore, the specific temporal coordination between Arp2/3 and VASP is crucial to the strongly accelerating protrusion. Notably, such molecular temporal coordination was reported to be involved in cell protrusion^12,13,42,43^. Particularly, PI3K has been known to increase after protrusion onset to stabilize nascent cell protrusion^42^. Taken together, our HACKS framework combined with pharmacological perturbations effectively demonstrated that heterogeneous edge movements could be deconvolved into variable protrusion phenotypes to reveal the underlying differential regulation of actin molecular dynamics. We also successfully demonstrated that we could monitor the changes in actin regulator dynamics induced by functional perturbation.

## Discussion

We have demonstrated that our computational framework HACKS could effectively deconvolve highly heterogeneous subcellular protrusion activities into distinct protrusion phenotypes, establish an association between each protrusion phenotype and the underlying differential actin regulator dynamics, and reveal specific phenotypes susceptible to pharmacological perturbations. Although previous studies have examined the spatiotemporal patterning of cell edge dynamics^11,44-46^, our study is the first to propose an effective framework to analyze the temporal heterogeneity in protrusion activities at the subcellular level. Using our framework, we have identified ‘fluctuating’, ‘periodic’, and ‘accelerating’ protrusion phenotypes of distinct temporal patterns in subcellular protrusion velocity. Based on the functional experiments using the pharmacological inhibitors, we further indentied ‘strongly’ and ‘weakly’ accelerating protrusions. Although previous studies also described persistent protrusion based on protrusion distance on a longer time scale^11,13,23,24^, our study is the first to further dissect protrusion phenotypes at a fine spatiotemporal scale and quantitatively characterize persistently ‘accelerating’ protrusion. Intriguingly, accelerating protrusion was later shown to be regulated by differential mechanisms, although they accounted for a minor portion of entire sampled protrusions. This finding indicates that identifying even a small subset of phenotypes is crucial to fully understand the biology underlying heterogeneous cellular behaviors. Furthermore, we also demonstrated that HACKS could be used for functional studies by pharmacological perturbations to achieve a more detailed mechanistic understanding of subcellular phenotypes.

We were also able to quantitatively measure how the underlying molecular dynamics are coordinated with protrusion phenotypes, thereby revealing the hidden variability of molecular regulatory mechanisms. Elucidating precise differential regulatory mechanisms related to protrusion heterogeneity has been difficult partly because it remains challenging to experimentally perturb a subset of molecules involved with specific subcellular phenotypes *in situ*. To address this challenge, our framework employed highly specific correlation and classification analyses. The result of the correlation analyses provided quantitative and detailed information about the differential coordination between molecular dynamics and the protrusion phenotype at the subcellular level. The classification analysis further established the association between each protrusion phenotype and the actin regulators by predicting the protrusion phenotypes using actin regulator dynamics.

We also demonstrated that we could deconvolve the heterogeneity of pharmacological responses of cellular protrusions using our HACKS framework by mapping protrusion time-series to two-dimensional phenotypic space using t-SNE and our time series clustering results. This approach revealed specific protrusion phenotypes which are most susceptible to pharmacological perturbations. Moreover, we were able to functionally validate our hypothesis drawn from the statistical analysis.

Our results suggest that the temporal coordination between Arp2/3 and VASP drives the acceleration of lamellipodial protrusion. To date, the Arp2/3 complex has been widely accepted as a master organizer of branched actin networks in lamellipodia that acts by nucleating actin filaments^29^, whereas VASP has been thought to be an elongator of actin filaments or anti-capper of the barbed ends^32,33,37,47^. In this study, we focused on the distinct recruitment dynamics of Arp3 and VASP identified in the accelerating protrusion phenotype (Cluster III). This suggested that Arp2/3-dependent actin nucleation provides a branched structural foundation for protrusion activity, and VASP-mediated actin elongation subsequently takes over to persistently accelerate protrusions. Our functional studies using CK666 and Cytochalasin D confirmed that this coordination is critical to strongly accelerating cell protrusion and the recruitment timing and duration of Arp3 and VASP is finely regulated to generate differential protrusion activities. Notably, VASP was reported to increase cell protrusion activities^26,27,39^, and has been implicated in cancer invasion and migration^26,48,49^. Thus, the coordination of Arp2/3 and VASP may regulate the plasticity of protrusion phenotypes, and the functional deregulation of the VASP or its isoforms in cancer may promote cellular migratory behaviors by promoting accelerating protrusion. Therefore, a mechanistic understanding of the connection between the specific protrusion and long-term migration may allow us to understand how distinctive migratory behavior arises among populations and shed light on how a subset of cancer cells acquire metastatic ability.

Lastly, our study is the first to quantitatively dissect the subcellular heterogeneity of dynamic cellular behaviors in unperturbed and perturbed cell states, which requires the deconvolution of temporal patterns in a highly fluctuating dynamic dataset. We do not consider HACKS to be limited to the analyses of subcellular protrusion heterogeneity: we anticipate that it can be expanded to study the morphodynamic heterogeneity of other types of cytoskeletal structures and membrane-bound organelles. Together, with the further development of unsupervised learning along with an increased repertoire of molecular dynamics, we expect our machine learning framework for live cell imaging data to accelerate the mechanistic understanding of heterogeneous cellular and subcellular behaviors.

**Supplementary Information** is available in the online version of this paper.

## Acknowledgements

We thank Seungeun Oh and Joseph Brazzo for the critical reading of the manuscript. We thank NVIDIA for providing us with TITAN X GPU cards (NVIDIA Hardware Grant Program) and Microsoft for providing us with Azure cloud computing resources (Microsoft Azure Research Award). This work is supported by the WPI Start-up Fund for new faculty, a generous gift by Boston Scientific, and NIH grant GM122012.

## Author Contributions

C. W. initiated the project, designed the algorithm of the time series clustering, performed the correlation analysis, and drug response analysis, and wrote the final version of the manuscript and supplement. H. C. performed the fluorescence live cell imaging, drug perturbation experiments, and wrote the final version of the manuscript and supplement; S. K. performed the classification analyses; A. D. performed cytoplasmic control experiments, N. L performed immunofluorescence experiments. D. K. and Y. B. contributed to the writing of the manuscript; K. L. coordinated the study and wrote the final version of the manuscript and supplement. All authors discussed the results of the study.

## Author Information

The authors declare no competing financial interests. Correspondence and requests for materials should be addressed to K.L.

## References

1 Small, J. V., Stradal, T., Vignal, E. & Rottner, K. The lamellipodium: where motility begins. Trends in cell biology 12, 112–120 (2002).

2 Pankov, R. et al. A Rac switch regulates random versus directionally persistent cell migration. The Journal of cell biology 170, 793–802, doi:10.1083/jcb.200503152 (2005).

3 Lauffenburger, D. A. & Horwitz, A. F. Cell migration: a physically integrated molecular process. Cell 84, 359–369 (1996).

4 Guirguis, R., Margulies, I., Taraboletti, G., Schiffmann, E. & Liotta, L. Cytokine-induced pseudopodial protrusion is coupled to tumour cell migration. Nature 329, 261–263, doi:10.1038/329261a0 (1987).

5 Morikawa, Y. et al. Actin cytoskeletal remodeling with protrusion formation is essential for heart regeneration in Hippo-deficient mice. Science signaling 8, ra41, doi:10.1126/scisignal.2005781 (2015).

6 Antonello, Z. A., Reiff, T., Ballesta-Illan, E. & Dominguez, M. Robust intestinal homeostasis relies on cellular plasticity in enteroblasts mediated by miR-8-Escargot switch. The EMBO journal 34, 2025–2041, doi:10.15252/embj.201591517 (2015).

7 Liu, Y. H. et al. Protrusion-localized STAT3 mRNA promotes metastasis of highly metastatic hepatocellular carcinoma cells in vitro. Acta pharmacologica Sinica 37, 805–813, doi:10.1038/aps.2015.166 (2016).

8 Taniuchi, K., Furihata, M., Hanazaki, K., Saito, M. & Saibara, T. IGF2BP3-mediated translation in cell protrusions promotes cell invasiveness and metastasis of pancreatic cancer. Oncotarget 5, 6832–6845, doi:10.18632/oncotarget.2257 (2014).

9 Ioannou, M. S. et al. DENND2B activates Rab13 at the leading edge of migrating cells and promotes metastatic behavior. The Journal of cell biology 208, 629–648, doi:10.1083/jcb.201407068 (2015).

10 Leithner, A. et al. Diversified actin protrusions promote environmental exploration but are dispensable for locomotion of leukocytes. Nat Cell Biol 18, 1253–1259, doi:10.1038/ncb3426 (2016).

11 Machacek, M. & Danuser, G. Morphodynamic profiling of protrusion phenotypes. Biophysical journal 90, 1439–1452, doi:10.1529/biophysj.105.070383 (2006).

12 Machacek, M. et al. Coordination of Rho GTPase activities during cell protrusion. Nature 461, 99–103, doi:10.1038/nature08242 (2009).

13 Lee, K. et al. Functional hierarchy of redundant actin assembly factors revealed by finegrained registration of intrinsic image fluctuations. Cell systems 1, 37–50, doi:10.1016/j.cels.2015.07.001 (2015).

14 Altschuler, S. J. & Wu, L. F. Cellular heterogeneity: do differences make a difference? Cell 141, 559–563, doi:10.1016/j.cell.2010.04.033 (2010).

15 Raj, A. & van Oudenaarden, A. Nature, nurture, or chance: stochastic gene expression and its consequences. Cell 135, 216–226, doi:10.1016/j.cell.2008.09.050 (2008).

16 Slack, M. D., Martinez, E. D., Wu, L. F. & Altschuler, S. J. Characterizing heterogeneous cellular responses to perturbations. Proceedings of the National Academy of Sciences of the United States of America 105, 19306–19311, doi:10.1073/pnas.0807038105 (2008).

17 Levine, J. H. et al. Data-Driven Phenotypic Dissection of AML Reveals Progenitor-like Cells that Correlate with Prognosis. Cell 162, 184–197, doi:10.1016/j.cell.2015.05.047 (2015).

18 Patel, A. P. et al. Single-cell RNA-seq highlights intratumoral heterogeneity in primary glioblastoma. Science 344, 1396–1401, doi:10.1126/science.1254257 (2014).

19 Shafqat-Abbasi, H. et al. An analysis toolbox to explore mesenchymal migration heterogeneity reveals adaptive switching between distinct modes. Elife 5, e11384, doi:10.7554/eLife.11384 (2016).

20 Sailem, H., Bousgouni, V., Cooper, S. & Bakal, C. Cross-talk between Rho and Rac GTPases drives deterministic exploration of cellular shape space and morphological heterogeneity. Open Biol 4, 130132, doi:10.1098/rsob.130132 (2014).

21 Huang, N. E. et al. The empirical mode decomposition and the Hilbert spectrum for nonlinear and non-stationary time series analysis. Proceedings of the Royal Society of London A: Mathematical, Physical and Engineering Sciences 454 (1998).

22 Keogh, E., Lin, J. & Fu, A. HOT SAX: Efficiently Finding the Most Unusual Time Series Subsequence. In Proc. of the 5th IEEE International Conference on Data Mining 226–233 (2005).

23 Rodriguez, A. & Laio, A. Machine learning. Clustering by fast search and find of density peaks. Science 344, 1492–1496, doi:10.1126/science.1242072 (2014).

24 Davies, D. L. & Bouldin, D. W. A cluster separation measure. IEEE transactions on pattern analysis and machine intelligence, 224–227 (1979).

25 Caliński, T. & Harabasz, J. A dendrite method for cluster analysis. Communications in Statistics-theory and Methods 3, 1–27 (1974).

26 Bae, Y. H. et al. Profilin1 regulates PI(3,4)P2 and lamellipodin accumulation at the leading edge thus influencing motility of MDA-MB-231 cells. Proceedings of the National Academy of Sciences of the United States of America 107, 21547–21552, doi:10.1073/pnas.1002309107 (2010).

27 Barnhart, E. L., Allard, J., Lou, S. S., Theriot, J. A. & Mogilner, A. Adhesion-Dependent Wave Generation in Crawling Cells. Curr Biol 27, 27–38, doi:10.1016/j.cub.2016.11.011 (2017).

28 Los, G. V. et al. HaloTag: a novel protein labeling technology for cell imaging and protein analysis. ACS Chem Biol 3, 373–382, doi:10.1021/cb800025k (2008).

29 Pollard, T. D. & Borisy, G. G. Cellular motility driven by assembly and disassembly of actin filaments. Cell 112, 453–465 (2003).

30 Benjamini, Y. & Hochberg, Y. Controlling the false discovery rate: a practical and powerful approach to multiple testing. Journal of the royal statistical society. Series B (Methodological), 289–300 (1995).

31 Rottner, K., Behrendt, B., Small, J. V. & Wehland, J. VASP dynamics during lamellipodia protrusion. Nat Cell Biol 1, 321–322, doi:10.1038/13040 (1999).

32 Barzik, M. et al. Ena/VASP proteins enhance actin polymerization in the presence of barbed end capping proteins. J Biol Chem 280, 28653–28662, doi:10.1074/jbc.M503957200 (2005).

33 Breitsprecher, D. et al. Clustering of VASP actively drives processive, WH2 domain-mediated actin filament elongation. The EMBO journal 27, 2943–2954, doi:10.1038/emboj.2008.211 (2008).

34 Hansen, S. D. & Mullins, R. D. Lamellipodin promotes actin assembly by clustering Ena/VASP proteins and tethering them to actin filaments. Elife 4, doi:10.7554/eLife.06585 (2015).

35 Machesky, L. M. et al. Scar, a WASp-related protein, activates nucleation of actin filaments by the Arp2/3 complex. Proceedings of the National Academy of Sciences of the United States of America 96, 3739–3744 (1999).

36 Nolen, B. J. et al. Characterization of two classes of small molecule inhibitors of Arp2/3 complex. Nature 460, 1031–1034, doi:10.1038/nature08231 (2009).

37 Rotty, J. D. et al. Profilin-1 serves as a gatekeeper for actin assembly by Arp2/3-dependent and-independent pathways. Dev Cell 32, 54–67, doi:10.1016/j.devcel.2014.10.026 (2015).

38 Bear, J. E. et al. Antagonism between Ena/VASP proteins and actin filament capping regulates fibroblast motility. Cell 109, 509–521 (2002).

39 Lacayo, C. I. et al. Emergence of large-scale cell morphology and movement from local actin filament growth dynamics. PLoS Biol 5, e233, doi:10.1371/journal.pbio.0050233 (2007).

40 Neel, N. F. et al. VASP is a CXCR2-interacting protein that regulates CXCR2-mediated polarization and chemotaxis. J Cell Sci 122, 1882–1894, doi:10.1242/jcs.039057 (2009).

41 Rosvall, M. & Bergstrom, C. T. Maps of random walks on complex networks reveal community structure. Proceedings of the National Academy of Sciences of the United States of America 105, 1118–1123, doi:10.1073/pnas.0706851105 (2008).

42 Welf, E. S., Ahmed, S., Johnson, H. E., Melvin, A. T. & Haugh, J. M. Migrating fibroblasts reorient directionality by a metastable, PI3K-dependent mechanism. The Journal of cell biology 197, 105–114, doi:10.1083/jcb.201108152 (2012).

43 Johnson, H. E. et al. F-actin bundles direct the initiation and orientation of lamellipodia through adhesion-based signaling. The Journal of cell biology 208, 443–455, doi:10.1083/jcb.201406102 (2015).

44 Martin, K. et al. Spatio-temporal co-ordination of RhoA, Rac1 and Cdc42 activation during prototypical edge protrusion and retraction dynamics. Sci Rep 6, 21901, doi:10.1038/srep21901 (2016).

45 Verkhovsky, A. B. The mechanisms of spatial and temporal patterning of cell-edge dynamics. Curr Opin Cell Biol 36, 113–121, doi:10.1016/j.ceb.2015.09.001 (2015).

46 Dobereiner, H. G. et al. Lateral membrane waves constitute a universal dynamic pattern of motile cells. Phys Rev Lett 97, 038102, doi:10.1103/PhysRevLett.97.038102 (2006).

47 Hansen, S. D. & Mullins, R. D. VASP is a processive actin polymerase that requires monomeric actin for barbed end association. The Journal of cell biology 191, 571–584, doi:10.1083/jcb.201003014 (2010).

48 Carmona, G. et al. Lamellipodin promotes invasive 3D cancer cell migration via regulated interactions with Ena/VASP and SCAR/WAVE. Oncogene 35, 5155–5169, doi:10.1038/onc.2016.47 (2016).

49 Philippar, U. et al. A Mena invasion isoform potentiates EGF-induced carcinoma cell invasion and metastasis. Dev Cell 15, 813–828, doi:10.1016/j.devcel.2008.09.003 (2008).

